# The role of habitat configuration in shaping animal population processes: a framework to generate quantitative predictions

**DOI:** 10.1101/2020.07.30.228205

**Authors:** Peng He, Pierre-Olivier Montiglio, Marius Somveille, Mauricio Cantor, Damien R. Farine

## Abstract

By shaping where individuals move, habitat configuration can fundamentally structure animal populations. Yet, we currently lack a framework for generating quantitative predictions about the role of habitat configuration in modulating population outcomes. For example, it is well known that the social structure of animal populations can shape spreading dynamics, but it remains underexplored to what extent such dynamics are determined by the underlying habitat configuration. To address this gap, we propose a framework and model inspired by studies using networks to characterize habitat connectivity. We first define animal habitat networks, explain how they can integrate information about the different configurational features of animals’ habitats, and highlight the need for a bottom-up generative model that can depict realistic variations in habitat structural connectivity. Second, we describe a model for simulating animal habitat networks (available in the R package *AnimalHabitatNetwork*), and demonstrate its ability to generate alternative habitat configurations based on empirical data, which forms the basis for exploring the consequences of alternative habitat structures. Finally, we use our framework to demonstrate how transmission properties, such as the spread of a pathogen, can be impacted by both local connectivity and landscape-level characteristics of the habitat. Our study highlights the importance of considering the underlying habitat configuration in studies linking social structure with population-level outcomes.

## Introduction

Animals rarely move unrestrictedly, as the physical habitat environments they depend on are often heterogeneous and uneven (Fahrig, 2007; Lovett et al., 2007; Kovalenko et al., 2012). By shaping individual movements, the physical configuration of habitats can have implications for population and community dynamics, including ecological interactions (Plitzko and Drossel, 2015; Jordano, 2016; Ryser et al., 2019), community structure (Altermatt and Holyoak, 2012; Henriques-Silva et al., 2013; Wilson et al., 2016), and speciation (Naka and Brumfield, 2018). Habitat configuration can also determine the rates of social interactions among conspecifics, thus shaping the social structure of populations (Emlen and Oring 1977; Gosling 1991; Leu et al., 2016; Farine and Sheldon, 2019; He et al., 2019). Ultimately, the physical configuration of habitats shapes the distributions of genes (Armansin et al., 2019), pathogens (Loehle, 1995; Altizer et al., 2003; Silk et al., 2019), and information (Laiolo and Tella, 2005, 2006; Aplin et al., 2015) in populations. Understanding the effects of habitat physical configuration on animal population and community dynamics is particularly important in a rapidly changing world, where natural populations increasingly face anthropogenic habitat changes. However, the role of habitat configuration in shaping population structure, and thus modulating population outcomes, remain underexplored.

How individual animals are socially structured has many consequences for populations (Keeling 1999; Aplin et al., 2012; Allen et al., 2017; Montiglio et al., 2018; Cantor et al., 2019). The best example for this perhaps comes from studies on pathogen transmission (Prado et al., 2009; Sah et al., 2018; Silk et al., 2019) describing how patterns of social or physical connections among individuals at local and global scales can impact the speed of transmission and the magnitude of disease outbreaks. Specifically, more clustered connections–where the number of shared social connections between individuals, or triads A↔B, B↔C, and A↔C, are more represented in the population – can increase the local spread (among immediate contacts) but decrease the speed and global reach of pathogen transmission (Read and Keeling, 2003; Keeling, 2005; Sah et al., 2018). However, to unravel the role of social structure in shaping ecological and evolutionary dynamics, we need to also understand the mechanisms that shape animal social structure. Alongside social decisions, features of the physical habitat environments can play a major role in shaping where animals move, who they (re-)encounter, and how often they interact with one-another (He et al., 2019). For example, a study in sleepy lizards (*Tiliqua rugosa*) found that habitats with more barriers increased the rates of encounters among individuals, increasing the density and clustering of the social networks (Leu et al., 2016), which may have implications for the spread of infectious pathogens (Tildesley et al., 2010; White et al., 2018). Early socioecological models have linked the spatiotemporal distribution of resources and risks to social behaviour (e.g. Wilson 1975; Van Schaik 1989), while more recent models have focused the behavioural mechanisms underlying social structure (Ilany and Akçay, 2016; Spiegel et al., 2016; Kappeler, 2017; Cantor and Farine, 2018; Farine, 2020). However, we also require quantitative tools to enable us to explicitly link configurational properties of habitats to social structures, and therefore generate testable hypotheses on the role of the physical habitat environments on socially-mediated population outcomes.

The features of animal habitats are typically multi-faceted–they can be described by the heterogeneity, sizes, abundance and spatial arrangements of habitat components (Tokeshi and Arakaki, 2012). For a given animal species, these features determine habitat structural connectivity, indicating where individuals can move and, thereby, the behaviours that they express. For example, Doherty et al. (2019) found that the shape of habitats, specifically whether habitats were wider (i.e. forming a rectangle) or thinner (forming a narrow strip), structured the movements of radio-tracked agamid lizards (*Pogona barbata*). Specifically, activity area and daily movement rates were lower among individuals inhabiting thinner habitats. The actual movements of animals are then the outcomes of both habitat structural connectivity and individuals’ behavioural decisions, which over time results in functional connectivity (see Box 1). An increasing number of studies have used networks for characterizing the structural or functional connectivity of animal habitats (Urban and Keitt, 2001; Bodin and Norberg, 2007; Fall et al., 2007; Minor and Urban, 2008; Urban et al., 2009; Lookingbill et al., 2010; Galpern et al., 2011; Alther and Altermatt, 2018; Marini et al., 2019; Poli et al., 2020). For example, Robertson et al. (2018) used long-term mark–resight data to construct networks that characterize the functional connectivity among habitat patches of snail kite (*Rostrhamus sociabilis plumbeus*) to evaluate the relative roles of among-patch movement and reproduction in modulating the effective connectivity of the species’ distribution range. In such networks, nodes often represent habitat or resource patches (Urban et al., 2009; Galpern et al., 2011), that is, areas crucial for survival and reproduction (Fahrig and Merriam, 1985) as opposed to the landscape matrix (Ziolkowska et al., 2014). How these patches are connected determines what these networks are depicting. Typically, functional connections are inferred from observed movement or from the flow of genetic material, while the structural connections are measures using Euclidean distances or least-cost paths determined by characteristics of the environment itself (reviewed in Galpern et al., 2011, see also Box 1).

### Box 1 Two distinctive approaches for describing habitat connectivity

#### Habitat structural (or physical) connectivity and habitat networks

Structural connectivity captures the physical features of habitat components and their spatial arrangement, which defines where animals of a given species can move (Baguette and Van Dyck, 2007). Habitat networks fundamentally describe the features of the physical habitat environments that determine habitat structural connectivity for a species, rather than the actual movements of animals through their physical habitat environments. For example, the features and spatial distribution of the chambers in an ant colony (Perna and Theraulaz, 2017) determine the structural connectivity of the colony, which can then structure the movements – and other processes, such as recruitment–of ants (Vaes et al., 2020). The actual movements of individuals can also be the products of other non-habitat components, such as the instant social environment the individuals are exposed to. Thus, in contrast to functional connectivity and movement networks (see below), structural connectivity describes the habitat components and their spatial arrangements for a given species–the habitat networks–that determine the movement potential of individuals (Fig. 2).

#### Habitat functional connectivity and movement networks

Animal movement networks characterize the observed movements of animals, with nodes anchored in physical landscapes and links quantifying the movement flows of animals among them (Pasquaretta et al., 2020). The resulting functional connectivity then aims to describe the extent to which habitats facilitates or impedes movements (Taylor et al., 1993; Goodwin and Fahrig, 2002). However, movement networks are the products of the spatial, social, and stochastic components that shape individual movements (Jacoby and Freeman, 2016), and habitat functional connectivity therefore reflects the interplay between individual behavioural decisions of animals and their physical habitat environments (Baguette and Van Dyck, 2007) as well as other underlying drivers of movement decisions, which can be multiple and complex (Nathan et al., 2008). For example, the movements of individuals from one patch to another could be affected by both the habitat features between these patches (i.e. permeability of landscape matrix), as well as the social (or biological) environment (Armansin et al., 2019) and an inherent drive to move in a specific direction (e.g. migration). Thus, functional connectivity inferred from observed movements may not completely estimate the underlying landscape features that facilitate or impede animal movements. The correlation between habitat structural connectivity and functional connectivity will further be influenced by the sampling effort – more samples (e.g. many individuals across many years) are likely to generate a stronger correlation. Popular approaches for characterizing functional connectivity include resistance surface modelling (Spear et al., 2015) and circuit theory (McRae et al., 2008). Recent insights on network-based habitat functional connectivity modelling are reviewed by Chubaty et al. (2020).

Empirically-driven network approaches give us insights into how the key configurational features of habitats can be integrated into a consistent network-based framework for modelling animal habitat configuration. While networks constructed following the approaches outlined above have been instrumental in our understanding of how habitat-mediated animal movements can affect population outcomes (Robertson et al., 2018) and community structures (Altermatt and Holyoak, 2012), they typically do not allow us to make broader, or more general, predictions on the consequences of habitat physical configuration for animal populations and communities. The key limitation is that empirical networks typically focus on specific habitats for given populations (Baranyi et al., 2011; Baguette et al., 2013; e.g. Fig. 1-a), therefore they cannot be applied to characterize habitats with various configurational features (e.g. landscape linearity, spatial scale, etc.), and inferences made based on these networks cannot be generalized. For example, while we can empirically build networks to explicitly characterize the structural connectivity of the Yangtze and the Rhine rivers, these networks would fail to represent the full range of structural connectivity that we observe in freshwater riverine habitats. We therefore need tools designed to specifically focus on characterizing how the key features of the habitat (e.g. the spatial layouts and physical connectivity of headwaters, mid-reaches and lower-reaches in riverine habitats) translate to population outcomes (e.g. Carraro et al., 2020).

**Fig. 1.**
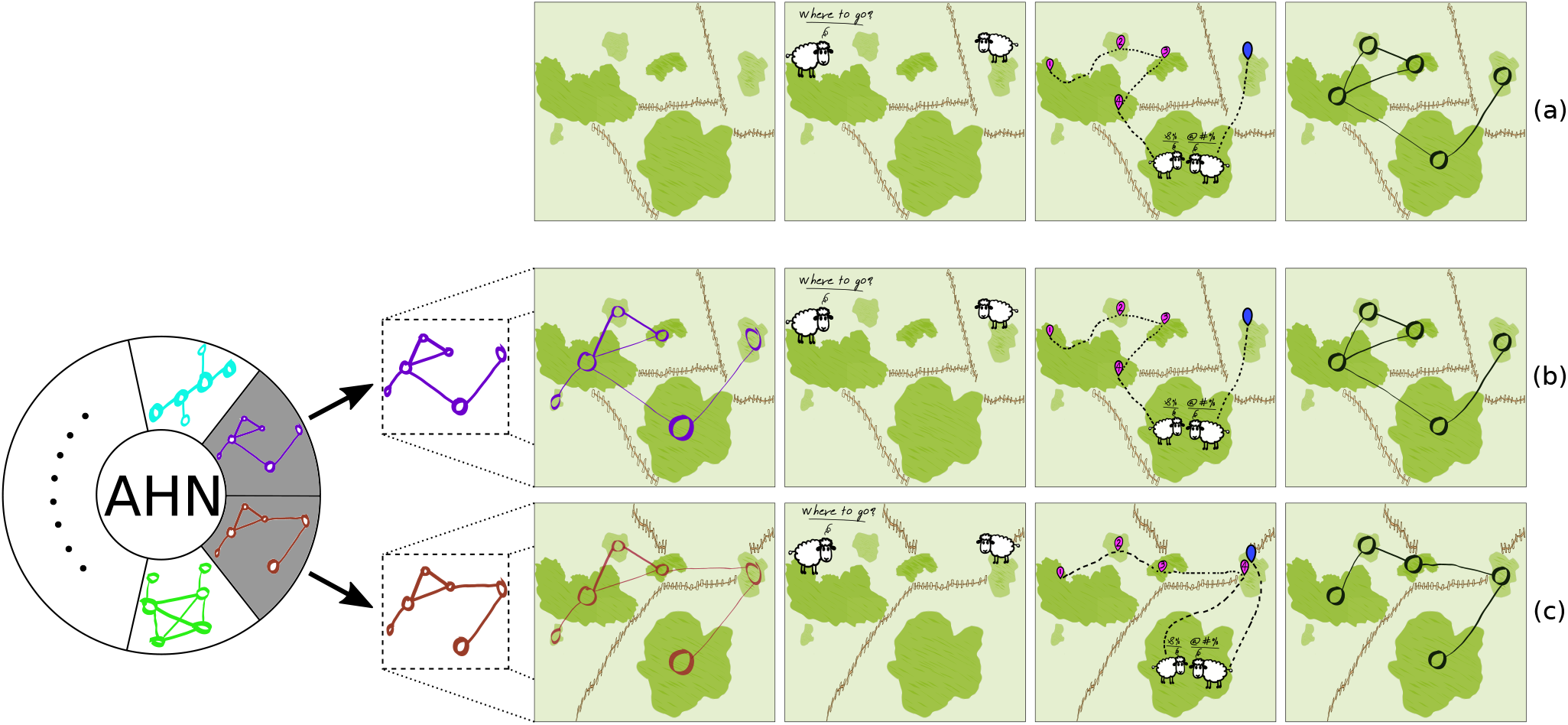
Two distinct approaches for understanding the role of habitat configuration in shaping animal population (or community) structures. (**a**) In most studies, animals are observed living and moving (e.g. via GPS tracking) within given time windows in specific habitats, from which characteristics of the structural or functional connectivity of the focal habitat area are inferred or modelled (e.g. by resistance surface modelling, network-based landscape connectivity modelling, or circuit theory, see Box 1). By contrast, (**b**) with a bottom-up approach, we can simulate networks to depict the physical configurations of specific habitats, and then model individual movements (or more complex behaviours) on these habitats, from which we can gain sights on how observed structures (e.g. patterns of movements and social interactions) emerged. With this approach, we can also (**c**) simulate habitat networks controlling for key parameters (e.g. network connectivity), thus producing alternative scenarios that can control (or not) for features that are hypothesized to play a major role in shaping biological processes in populations. Here we illustrate two simulated networks, one of which (**b**) can exactly depict the configuration of the given habitat (**a**), while the other depicts a habitat that maintains some characteristics (e.g. the same distributions and sizes of habitat patches, represented by nodes) as the given habitat (**a**), but provides alternative patterns of structural connectivity (by randomizing the spatial distribution of movement barriers that determine which patches are connected).

Making generalizable predictions about the consequences of habitat configuration requires being able to generate a range of alternative networks depicting a plausible diversity of habitat configurations, while at the same time being able to maintain certain key aspects of configurational features (e.g. Fig. 1-b, 1-c). One way to overcome the limitation of empirical networks is to simulate networks using a generative model. Generative network models have been instrumental to our understanding of the importance and consequences of network structure for a range of phenomena (Granovetter, 1973; Watts and Strogatz, 1998), but existing models currently cannot capture key structural properties in the connectivity of animal habitats (but see Carraro et al., 2020 for a recent solution for riverine networks). For example, animal habitat network connectivity is inherently spatially-dependent–it inherits many of its structural properties from the spatial distribution of habitat components, the features of landscape matrix, and the geometry of the landscape. General models of random networks (e.g. Erdős and Rényi, 1960; Barabási et al., 2001) do not consider such spatial properties, therefore creating network that intrinsically differ in patterns of structural connectivity compared to animal habitat networks. One such model of random networks incorporating spatial components is the random geometric network model (Dall and Christensen, 2002; Penrose, 2003). The nodes in geometric networks are anchored in space, while links are defined depending on the locations of nodes. That is, nodes are connected whenever their Euclidean distance is below a given threshold. Thus, the geometric model generates spatial networks in the sense that links depend on spatial locations of nodes (Barthélemy, 2011), capturing general properties of habitat networks, and thereby providing an ideal starting point for generating simulated habitat networks. However, current limitations of the random geometric network model include having a fixed spatial extent (i.e. a square) and a fixed threshold for connecting habitat components. In nature, the geometry of habitats and landscapes, and the properties of their connectivity, can vary remarkably. For example, the spatial distance between two patches does not exclusively dictate their connectivity–barriers such as waterscapes can restrict movement of a terrestrial animal between two patches in close proximity, while patches that are far apart can be connected. Several studies (e.g. Strandburg-Peshkin et al., 2017; Green et al., 2020) have demonstrated that primates can use roads to efficiently move between distant areas of their home ranges.

Here, we address the need for a generative model of animal habitat networks by extending the random geometric network model. We first define animal habitat networks and outline the key configurational features of animal habitats that can be captured by such networks. Next, we describe a modelling framework (available in the R package *Animal Habitat Network*) for simulating animal habitat networks that is explicitly tailored to depict the diverse physical configurations of animal habitats. We show that our network simulation algorithm can be tuned to efficiently capture the structural connectivity of real habitats. Doing so is important, as making predictions requires producing realistic scenarios as well as contrasting these against alternative scenarios. Finally, we illustrate how our framework can be used to simulate animal habitat networks with varying patterns of connectivity to investigate the implications of habitat configuration for populations by embedding a Susceptible-Infected-Recovered (SIR) epidemic model in simulated habitat networks. This can provide new insights on the link between habitat configuration and population-level outcomes. Taken together, our findings highlight how the application of an explicit and quantitative framework to simulate habitat networks can help us gain a better mechanistic understanding of the role of habitat configuration in shaping the dynamics of ecological, evolutionary processes and their conservation implications.

### Defining animal habitat networks

Animal habitats are defined by taking both the species-level properties (such as locomotion and space use characteristics of the focal species) and environmental features into account. We consider the five fundamental dimensions proposed for characterizing habitat physical configuration (Tokeshi and Arakaki, 2012) to define animal habitat networks (Fig. 2) as network-based explicit depictions of 1) *the spatial organization* and 2) *the physical attributes* of habitat components at given spatial scales, and 3) *the structural connectivity* (Box 1) indicating where animals of a given species can move. In habitat networks, nodes with attributes characterize the physical features of habitat components (e.g. location, size, quality, physical composition). The presence of a link between two nodes indicates that animals of a given species are physically able to move between these two habitat components (e.g. due to a physical distance compatible with the species mobility, and/or the absence of physical barriers). The weight of a link can characterize the spatial proximity or permeability of the non-habitat matrix between these two components (Fig. 2). In this way, the statistical properties of link weights in a habitat network mirror expected movement propensity among habitat components due to habitat physical features and species-level movement characteristics, rather than the realised movements of individuals among habitat components (i.e. functional connectivity, Taylor et al., 1993). Therefore, animal habitat networks (Fig. 2) are distinct from movement networks (Box 1), where links characterize the actual movements of animals among habitat components (Jacoby and Freeman, 2016; Pasquaretta et al., 2020). Animal habitat networks provide the fundamental physical templates for where animals can move, while the actual movements of individuals can also be the products of other drivers, such as the social environment that shape individuals’ movement decisions (Couzin et al., 2005; Strandburg-Peshkin et al., 2015). Our definition of animal habitat networks highlights the fundamental bottom-up role that habitat physical configuration, in and of itself, has in structuring animal movements and the subsequent processes in populations or communities.

**Fig. 2.**
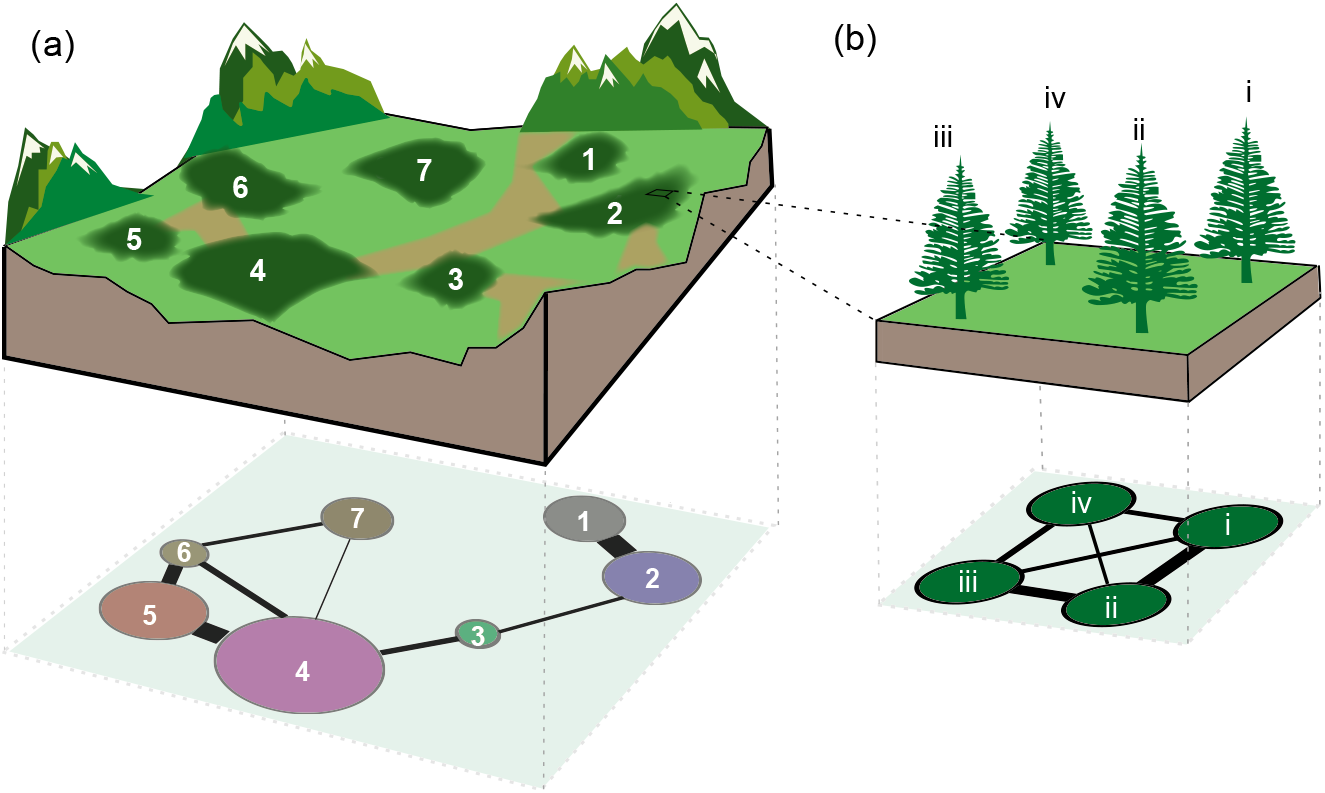
Networks explicitly characterizing the physical configurations of animal habitats. We illustrate how five dimensions for assessing habitat configuration proposed by Tokeshi and Arakaki (2012) can be integrated and applied to construct animal habitat networks. These dimensions are 1) *spatial scale* (spatial resolution and extent), 2) *composition diversity* (heterogeneity), 3) *size* (area), 4) *abundance or density* (number of discrete habitat units per area), and 5) *spatial arrangement* (distribution) of habitat components. (**a**) A hypothetical land-scape composed by forest fragments (numbered components) within a heterogeneous matrix with potential movement corridors (light green, which account for the presences of links between nodes) and physical barriers (light brown, which account for the absence of links between nodes). The physical features and spatial organization of the habitat components can be represented by a connected network at a large spatial scale, with a high composition diversity (fragments of different tree species), different habitat sizes (small and large fragments), high abundance (7 fragments), and heterogeneous spatial arrangement (fragments unequally distributed and connected by movement corridors across the landscape). (**b**) The physical features and spatial arrangement of habitat components can be characterized at different spatial scales. Here, part of the forest (fragment 2) can be represented by a connected network at a finer spatial scale (e.g. trees as habitat components), with a low composition diversity (the same tree species), small habitat size (single trees), low abundance of components (4 trees), and uniform spatial arrangement. In the two habitat networks, the compositional diversity (or quality) and size (or carrying capacity) of habitat components are characterized by node attributes (colours and sizes), the abundance by the number of nodes in the networks, and the spatial arrangement by the patterns of connectivity and the distribution of link weights (both as a function of the Euclidean distances between habitat components).

### AHN: a multi-dimensional framework for modelling animal habitat configuration

#### The model

We propose a general and spatially-explicit modelling framework for plausible animal habitat networks (hereafter the ‘AHN’) that can depict the diverse configurational features of animal habitats. The model contains eight parameters (Table 1) explicitly encoding five fundamental dimensions characterizing animal habitat physical configuration (Tokeshi and Arakaki, 2012) and species-level movement characteristics. We define the model within a 2-dimensional Euclidean space, by conceiving a planar rectangular landscape of an area *A* and a side length *L*. The spatial coordinates ***x*** and ***y*** of *N* habitat components are then drawn from the intervals [0, *L*] and [0, *A*/*L*] respectively, with their distributions determined by probability distributions of interest (e.g. the uniform distribution in Fig. 3-i). Alternatively, the spatial coordinates can be specified in the model. In this way, the AHN can depict landscapes with variable overall landscape shapes, or based on the geometric features of empirical animal habitats. Furthermore, with a given landscape, the model can explicitly capture the number (or density) of habitat components at the given spatial scale. Finally, in the model, the compositional diversity (i.e. heterogeneity) and size (or other physical properties) of habitat components can be included as node attributes, in vectors ***U*** and ***V*** respectively, the values of which can be quantitative or qualitative, and can be determined specifically or drawn at random, depending on the hypothesis of interest.

**Fig. 3.**
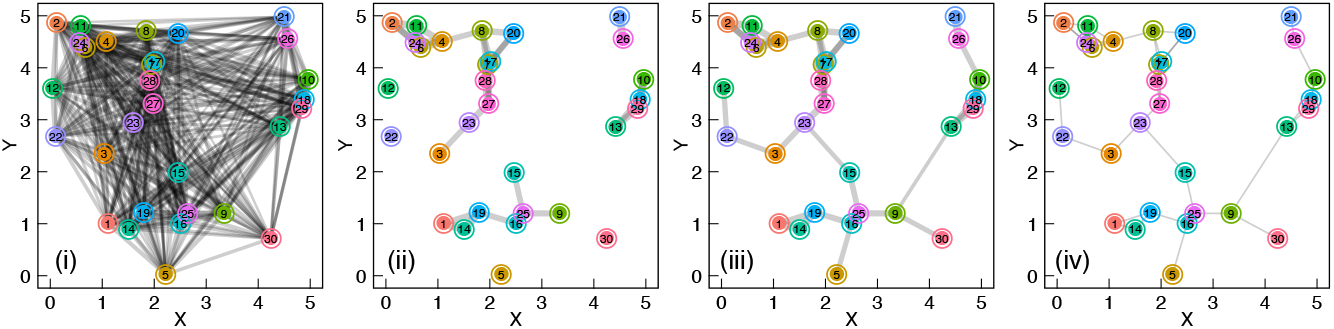
The workflow of the AHN model for generating animal habitat networks. First, (**i**) the algorithm constructs a fully connected and weighted habitat network. Here, numbered nodes represent 30 habitat components colour-coded by their attributes (such as their sizes, quality or compositions, with continuous or discrete colour palette) and connected by links whose thicknesses indicate the strength of the spatial relationship between the two habitat components, and is determined by the spatial positions of the nodes. The network is defined in a conceived 2-dimensional landscape in which the *x* and *y* axes indicate the spatial extents of the landscape (here the aspect ratio is 1, i.e. *A* = *L*^2^, but the model allows any *x* and *y* extents for capturing the diverse landscape geometry), therefore it inherits spatial properties of the landscape. Next, (**ii**) the algorithm removes the link between node *i* and *j*(*i* ≠ *j*) from the network with probability *P*(*D_ij_*); in this example it results in a disconnected habitat network. Then, (**iii**) the (disconnected) network components are rewired with minimal number of links. Finally, (**iv**) the habitat network can be transformed to unweighted, if so desired.

**Table 1.**
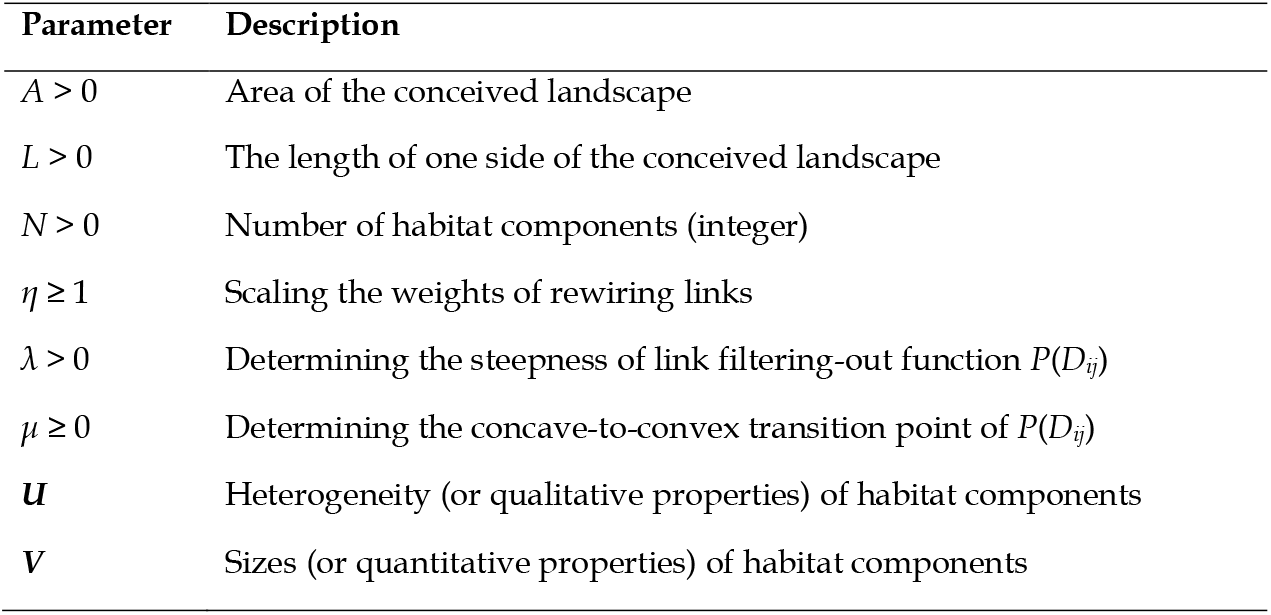
Parameters of the AHN model for depicting habitat physical configuration.

Once the layout of habitat components is defined, the AHN model can then generate links to characterize the structural connectivity between habitat components for a given species. Links can be weighted or unweighted, and are non-directed (the model can easily be extended to have directed links, for example if there is a gradient–such as altitude – a physical features of habitat that might favour animal movements in one direction more than the other). The model starts by allocating the link weights between node *i* and node *j* (*i* ≠ *j*) using the power function *W*(*D_ij_*) = *D_ij_*^-1^, where *D_ij_* is the Euclidean distance, a primary metric characterizing habitat structural connectivity (Poli et al., 2020), between habitat components that node *i* and node *j* are referring to (Fig. 3-i). With this approach, the network starts by being fully connected, with each weighted link indicating the spatial proximity or relation between pairs of habitat components. In natural environments, however, animals of a species at a given site usually cannot move directly to all other sites, because their movements are often restricted by the features of landscape matrix, such as physical barriers. Thus, we use the sigmoidal function *P*(*D_ij_*) = [1 + exp(-λ(*D_ij_* - *μ*))]^-1^ to determine a threshold probability for *filtering out* links between node *i* and *j* from the initial complete network. The dependence of the probability on *D_ij_* assumes that it is less likely that there exists a direct movement corridor between two habitat components when they are much further away (i.e. it is more likely that there will present inhospitable features to movements in the landscape matrix between the two distant sites). The probabilistic nature of this function captures the stochasticity that exists in the relationships between distance and connectivity–that is, most nearby components are connected, but some have barriers between them. The parameter *λ* determines the steepness of the thresholding curve (i.e. the extent to which distance determines the presence or absence of links), while *μ* ≥ 0 determines the critical distance *D_ij_* at which the curve transits from concave (more likely to be linked) to convex (less likely to be linked). We define *λ* > 0 so *P*(*D_ij_*) consistently increases over *D_ij_* (Fig. A1). This function enables us to generate a wide spectrum of curves to cover the diverse and evolving relationships between *D_ij_* and *P*(*D_ij_*) by tuning *λ* and *μ* (Fig. A1-A3)–which depict the propensity for physical features that inhibit individual movements (e.g. barriers) to exist in the landscape (*λ*) and the species-level movement characteristics that determine individuals’ ability to exploit existing physical connectivity (*μ*).

With such an approach, the framework allows us to simulate networks that can capture the diverse patterns of structural connectivity of animal habitats as found in nature. For example, with a given set of spatially-referenced nodes, *P*(*D_ij_*) allows us to simulate variable patterns of network connectivity, which can change properties of the network such as its clustering coefficient. *P*(*D_ij_*) can also determine the presence of links in both probabilistic and deterministic ways, which not only makes the framework general, but also enriches our ability to encode the diverse physical features between habitat components, such as the variation in the quality of non-habitat matrix (in terms of their effects on animal movements). For instance, in an extreme case when *μ* → 0 and *λ* → ∞, the link removal function becomes more deterministic (i.e. *P*(*D_ij_*) → 1), and the model can then generate networks that approximate planar networks (McDiarmid et al., 2005), which have previously been used to model landscape functional connectivity (Chubaty et al., 2020).

In some cases, *P*(*D_ij_*) fragments the network into (disconnected) network components (e.g. Fig. 3-ii). The more *μ* approaches 0, the more links will be filtered out, and the more likely it is for the network to be disconnected (Fig. A3). In such cases, the disconnected networks would represent habitats containing isolated clusters of habitat components between which animals cannot physically penetrate. From a modelling perspective, it is often preferable (at least initially) to consider one habitat as a connected network (i.e. a single network component, denoting a habitat or a section of a larger but fragmented habitat). Thus, we use a step-wise approach to rewire network components, with the two spatially closest nodes from each of the two network components being wired each time until the network has no disconnected network components, and the algorithm uses the minimum number of rewiring links for doing this (Fig. 3-iii). After rewiring, the weights of links rewiring network components are determined by the function G(*D_aiβj_, η*) = (*D_aiβj_*)^−*η*^, where *D_aiβj_* is the shortest Euclidean distance between node *i* from the network component *a* and node *j* from the network component *β*(*a* ≠ *β*), and *η* ≥ 1 mediates the strength of the connectivity between the two clusters of habitat components (Fig. A4). We provide the implemented algorithm in the function *ahn_gen()* in the R package *Animal Habitat Network* (version 0.1.0, He and Farine, 2019).

### Demonstrating the capability of the AHN model in simulating habitat structural connectivity

To test the capability of the AHN model in depicting the structural connectivity of animal habitats, we compared the topological properties of networks generated by the model using a given parameter space with those of empirical habitat networks characterizing habitat structural connectivity by Friesen et al. (2019). First, we extracted the largest network component from each of the 62 empirical habitat networks, and kept 58 of them for benchmarking (the two largest were omitted due to computation limit and the two simplest were excluded). Each of these extracted networks is connected, denoting a habitat or a part of a larger habitat where animals can physically move from a given habitat component to any other one (i.e. the habitat can be functionally connected–biological processes such as information or genes flows are possible among habitat components). Next, we simulated random habitat networks with the AHN model and identified those sets of parameters under which the corresponding output habitat networks best approximated the (average) clustering coefficient, modularity, and diameter of each of these empirical networks, respectively. We considered the parameter space *A* = 25, *L* ∈ {5, 10, 15, 20, 25, 30}, *μ* ∈ {0.1, 2, 5, 7, 10}, *λ* ∈ {0.001, 0.1, 0.15, 0.35, 0.4, 0.75, 1.25, 5, 30} across all empirical benchmark networks, while keeping *N* identical to the number of nodes of the corresponding empirical habitat network. This parameter space was determined by considering the effects of each parameter on the resulting network structures (Fig. A1-A7). In total, for each metric of each empirical network, we generated 270 (i.e. size of the parameter space) random habitat networks, from which the bestfitting set of parameters was determined. Then, we simulated 15 habitat networks with each of these sets of parameters as replicates, and evaluated the extent to which each of these metrics of each replicate deviated from that of the corresponding empirical network. These networks confirm that our model can generate simulated networks that capture the key structural properties of given habitats (Fig. 4), thereby forming the basis for subsequently exploring on how population outcomes (structure and/or processes) might change under alternative scenarios (e.g. by changing a key parameter from a given landscape, such as increasing or decreasing connectivity by tuning *μ*). All network computations were done in R (version 3.6.1, R Development Core Team, 2019) with the *igraph* library (version 1.2.5, Csardi and Nepusz, 2006).

**Fig. 4.**
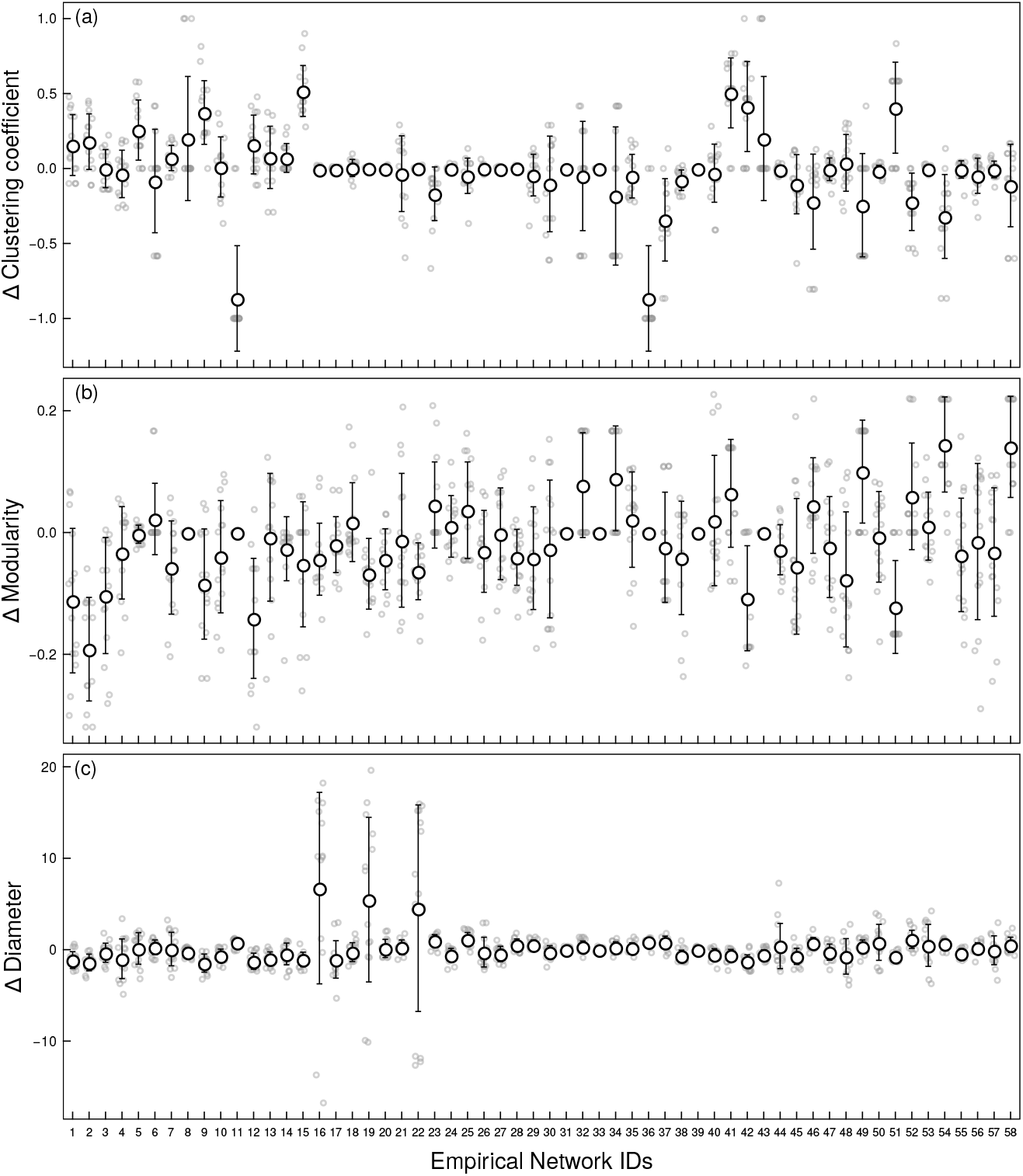
The AHN model can simulate spatially-explicit networks to characterize habitat structural connectivity. Each grey circle denotes the difference in each of the three metrics (y-axes) between each of the 15 replicated random habitat networks generated by the AHN model with each set of best-fitting parameters identified from the given parameter space and the corresponding empirical network; black circles and bars characterize the means and the standard deviations.

### Key research questions

The Animal Habitat Network model can be used to address a range of different topics and research questions. Here we provide some examples of how our framework can be used to directly extend existing studies. We identify three key research areas where our framework can be used to directly address outstanding gaps in knowledge.

### What are the consequences of habitat changes for animal population processes?

Animal habitats are changing under natural and anthropogenic drivers, typically characterized by changes in the spatial distribution of habitat components (e.g. food and shelters), changes in their structural connectivity (e.g. through fragmentation or reforestation), and/or changes in the physical attributes of the habitat components themselves (e.g. the amount of resources in each habitat patch). These changes can then reshape the movements of animals, which can subsequently affect the patterns of biological or ecological interactions (e.g. inter-individual social structure, predator-prey interactions), or even impose evolutionary pressures on impacted species (Kokko and Sutherland, 2001; Banks et al., 2011). A number of studies have shown that the spatial distributions of food resources or habitat fragmentation can shape the spatial organization of individuals (Mourier et al., 2012; Jacobson et al., 2015), with consequences on the evolution of their social or mating systems (Emlen and Oring, 1977; Van Schaik 1989; Banks et al., 2007; Tuomainen and Candolin, 2011). For example, Banks et al. (2011) empirically explored the relationship between the patterns of den-sharing interactions among hollow-dependent Australian mountain brushtail possums and the spatial variation in hollow tree availability, and noticed a behavioural switch from kin avoidance to kin preference in den sharing when hollow tree availability decreases, highlighting the important role of habitat change in driving individuals’ social behaviours as responses. In another example, Bain et al. (2014) examined the effects of habitat configuration on the frequency of extra-pair paternity (EPP) in cooperatively breeding superb fairy-wren (*Malurus cyaneus*) by linking spatial arrangements of their territories to the frequency of EPP, and found that the frequency of extra-group paternity (EGP) among groups in linear strips of vegetation was lower than those in more clustered territories in continuous habitats, highlighting the role of habitat spatial configuration in influencing the rates of EGP and the potential consequences of anthropogenic habitat change for mating systems.

Our network-based modelling framework can be used to depict multiple yet diverse configurational properties of animal habitats, thereby providing the starting point for explicitly modelling habitat change and predicting the population outcomes. Moreover, understanding the consequences of habitat fragmentation for populations is one of the central topics in conservation biology (Fischer and Lindenmayer, 2007; Haddad et al., 2015), while the impacts on social processes of conservation efforts aimed at reducing habitat fragmentation are almost completely unexplored. With our framework, one can simulate network scenarios that realistically map the potential trajectories of habitat change (e.g. by parameter optimization and/or network manipulation), and generate predictions on the potential consequences for a population.

### How is habitat connectivity shaped by landscape and species properties?

The patterns of connectivity that form animals’ habitats are shaped by a range of properties. Some of these are biological–such as the movement capacity of the animal species. Many of these properties are abiotic, such as the climatic conditions that determine the composition of habitat patches (e.g. the assemblage of plants, or coral, species in a patch) and geological features that determine the shape of the landscape (e.g. the long and narrow valleys created by a mountain range vs. an open plain). A key question is, therefore, whether certain types of landscapes consistently generate networks with different properties. For example, it is likely that riverine habitats, or habitats in valleys, will have a larger network diameter (i.e. a longer minimum movement path between the most distant nodes) than habitats that are not restricted by geometric features of landscapes. Using our framework, it will be possible to explore, and develop an understanding, of the relative roles of species-level properties (e.g. locomotion characteristics and mobility) and landscape properties (e.g. linearity) in shaping characteristic properties of habitat networks (e.g. network clustering).

### How different do we expect population social structures to be in different landscapes?

Studies have revealed that animal population social structures often exhibit notable variations (Whitehead and Kahn, 1992; Mori and Saito, 2005; Nandini et al., 2017; Prehn et al., 2019). When habitats vary in their physical configurations, we would expect the social structures of populations in these habitats to exhibit variations (even for the same species), and this is indicated by empirical evidence. For example, Farine and Sheldon (2019) showed that the social network structure (at the network community level) of a woodland bird community observed in the Wytham Woods in the UK, remained consistent across 4 winters, despite the high turnover rate of individuals within the communities. This study suggests that the predictability of habitat configuration for the emergent social network structures. Our framework can be tuned to depict animal habitats with distinct configurational features, therefore providing a theoretical tool to examine how much variations in population social structures observed from one habitat to another might be explained by habitat physical configurations (i.e. the extent to which animal habitat networks account for the variations in animal social networks). Figure 5 (from our example below) highlights how different aspects of habitat configuration (such as the aspect ratio and the tendency for distant patches to be connected versus not) can interact with each other to shape the resulting structural features of the habitat networks.

**Fig. 5.**
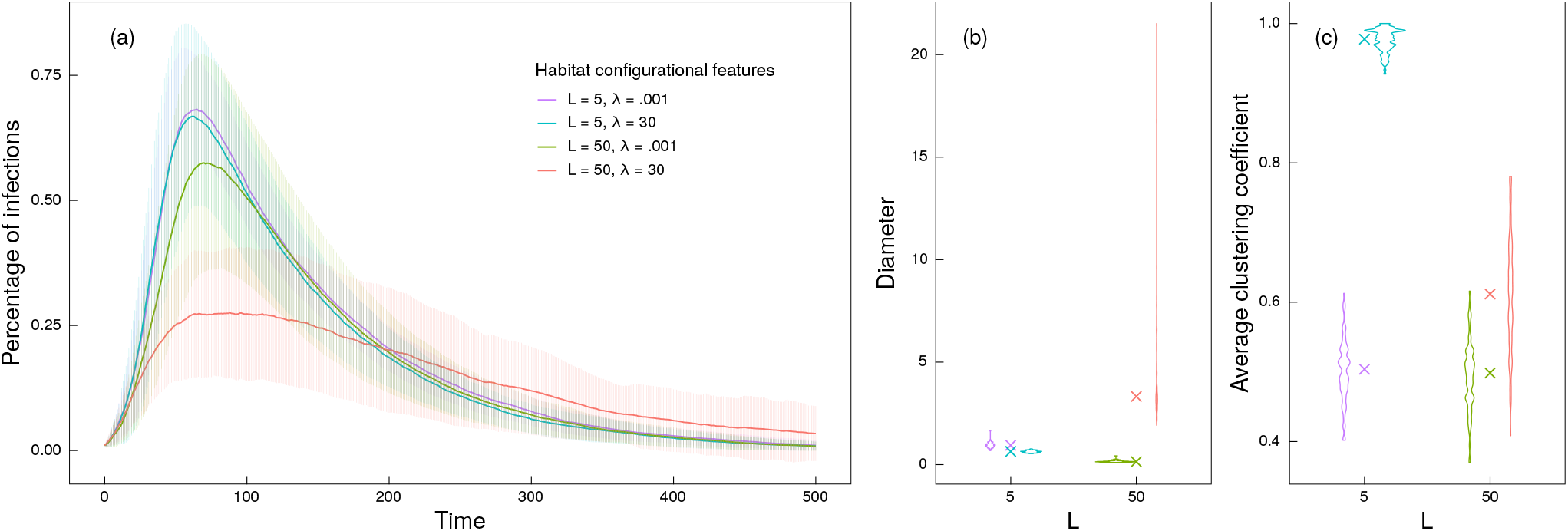
The application of the AHN model for understanding the role of habitat shape and structural connectivity in mediating pathogen transmission dynamics in habitat-structured animal populations. (**a**) The transmission of pathogens in populations is dependent on both the landscape shape, depicted by *L* where a larger value represents landscapes with a larger aspect ratio, and the extent to which the physical connections between patches to be determined by the distance between them, depicted by *λ* where a lower value corresponds to a weaker relationship between the distance and the tendency of being physically connected. Simulations show that the transmission dynamics, when individual mobility is at a medium level (1 - *p_s_* = 0.5) under an infection rate of *π* = 0.05 and a recovery rate of *ρ* = 0.01, are impacted by habitat shape and structural connectivity. Specifically, habitats with a larger aspect ratio (a larger *L* value) and with their structural connectivity determined by a stronger relationship between the distance and the probability of being connected (a larger *λ* value) have the smallest disease outbreaks (each curve indicates the mean percentage of infected individuals in a population of 100 individuals moving on simulated habitat networks comprising 20 nodes, over 500 timesteps, with bars indicating the standard deviations across 100 replications). The interaction between landscape shape and the degree to which between-patch physical connectivity is determined by distance affects properties of the habitat networks, such as their diameter (**b**) and their average clustering coefficients (**c**).

### Illustrating the framework: how do habitat networks shape pathogen transmissions?

We illustrate the application of our framework in understanding the link between animal habitat network structure and population outcomes in the context of pathogen transmission in a population of 100 individuals (i.e. with no birth, death, emigration and immigration) moving on habitat networks of size 20 simulated by the AHN model. We considered the simplest case where habitat patches are randomly positioned (with their *x* and *y* coordinates drawn from uniform distributions) in landscapes with the same area (i.e. *A* = 25) but varying aspect ratios–with *L* = 5 producing square-shaped landscapes (e.g. resembling a forest on an open plain) and *L* = 50 producing landscapes with a large aspect ratio (e.g. resembling a forest in a narrow valley). Within these landscapes, we considered two scenarios of habitat structural connectivity, defined by *λ* = 0.001 and *λ* = 30. These values depict two distinct relationships between the distance and the probability of there being a physical feature (e.g. physical barrier) restricting movement between two patches, whereby *λ* = 0.001 represents a weaker relationship between distance and the presence (or absence) of such feature, and *λ* = 30 represents a stronger relationship. We maintained other parameters at constant values (*μ* = 5 and *η* = 1).

We initiated simulations by randomly allocating individuals to the nodes of each habitat networks, and modelled individual movements and transmission dynamics for 500 timesteps. When modelling individual movements, we defined the probability for an individual to stay at the current node *i* at each timestep consistently as *p_s_* = 0.5, and the probability of moving from node *i* to *j* as (1 - *p_s_*) × *w_ij_* / Σ*_j_w_ij_* (where *w_ij_* is the weight of the link between node *i* and *j*). We simulated pathogen transmissions in the population using an SIR epidemic model (Kermack and McKendrick, 1927; Keeling and Eames, 2005), where each individual is either susceptible (S), infectious (I) or recovered/removed (R). We randomly set one individual (1% of the population) as infectious, and at each timestep, simulated infected individual infecting each of its susceptible neighbours in the same patch (if any) with probability *π* = 0.05. Individuals recovered and acquires immunity with the probability *ρ* = 0.01. We tracked the percentages of infected individuals in the simulated populations over time, and compared the mean percentages of infections observed from 100 replications.

We show that the landscape properties play a role in mediating the pathogen transmission dynamics. Importantly, large- (landscape geometry) and local- (the propensity for nearby patches to be more connected) scale characteristics work together to shape the predicted transmission dynamics of a simulated pathogen. A weaker relationship between inter-patch physical connectivity and distance (as modelled by a lower *λ* value) increases the scale of global disease outbreaks, but this effect is most strongly realised in landscapes with a larger aspect ratio (Fig. 5-a), such as a forest habitat in a narrow valley. Our results relate to existing literature linking social (or contact) network structures to patterns of disease transmissions–the stronger tendency for distant patches to be physically connected (i.e. *λ* = 0.001) and squarer landscapes (i.e. *L* = 5) typically decrease the path length, or diameter (Fig. 5-b), and clustering (Fig. 5-c) of the habitat networks, and correspondingly, increase the pathogen outbreak size in simulated populations. Our results complement recent work demonstrating that the fragmentation of animal habitats can impact the transmission dynamics of pathogens (Silk et al., 2019), extending it by showing that the shape as well as the internal connectivity of a habitat is important.

## Discussion

We present a multi-dimensional framework for simulating networks that can realistically capture the diverse physical configurations of animal habitats. Our model provides a tool to develop a more mechanistic understanding of the role of habitat configuration in modulating population processes and outcomes. Such modulating effects are likely to be widespread – for example we have demonstrated that the structure of the habitat network can have consequences on pathogen transmission, in line with predictions from studies of social networks. Developing such mechanistic knowledge is critical as natural animal populations face increasingly rapid changes in their habitats, which have ecological and evolutionary consequences. For instance, habitat change can affect the magnitude of competition (Calizza et al., 2017) and the spread dynamics of pathogens (Bloomfield et al., 2020), information (Betts et al., 2008), or genes (Keller and Largiader, 2003).

When modelling animal habitat physical configuration, the structural connectivity of habitats is key. Habitat networks are spatial networks, and their topological properties are inherently shaped by the spatial distributions of habitat components and the landscape features (such as the matrix quality and geometry of landscapes). From a theoretical perspective, existing models of generic and geometric random networks do not currently capture the rich structural connectivity of real habitats, as they are not tailored to depict the structural connectivity of animal habitats observed in nature. By taking a bottom-up approach to capturing habitat configurations, our model can be tuned to approximate the structural connectivity of specific habitat configurations. From these, researchers can produce a range of alternative, yet realistic, scenarios to explore the consequences of different features of the habitat on population processes, such as changes in the spatial arrangements of resource patches and physical barriers to movement.

The fundamental role of the physical environment on animal populations makes the evaluation of the consequences of habitat configuration relevant to both theorists and empiricists. If we do not explicitly consider habitat configuration, we risk missing the importance of its contribution to biological processes. For example, simulating social networks of large populations without considering spatial dependencies could produce networks that are more connected than they should (i.e. without considering the spatial constraints on social interactions). Doing so can misrepresent the biological processes that network structure shapes, such as the transmissions of pathogens (White et al., 2018; Wilkinson et al., 2018), information (Aplin et al., 2015), genes (Vähä et al., 2007). By contrast, current geometric network models, which are spatially-dependent, may largely overestimate the spatial clustering of habitat components because it does not allow for rare long-distance connections or missing connections among close patches by forcing all closely-located components to be connected. The need for models tailored to simulate habitat networks has been highlighted by recent studies that modelled specific habitat scenarios. For example, Carraro et al. (2020) proposed a toolkit for generating networks to capture the topological features of real riverine habitats to understand their role in shaping the key processes in freshwater ecology and evolution. The AHN model herein proposed is a more general and flexible framework for depicting habitat structural connectivity. Notably, when simulating networks, the AHN model allows any spatial distributions of habitat components in any landscape (i.e. by tuning *A* and/or *L*), and provides a cluster of probability curves (i.e. by tuning the *λ* and *μ* in the *P*(*D_ij_*) to model the diverse structural connectivity among habitat components. Also, the AHN can generate alternative representations of the structural connectivity of given habitats, and allows control over the deviations of alternative scenarios from the specific habitats that can be used to generate scenarios. Thus, our proposed model can generate realistic habitat scenarios that are biologically meaningful.

There are many useful applications in generating realistic animal habitat scenarios. For example, there is growing interest in understanding the interplay between habitat physical configurations and individuals’ behaviours to predict the persistence of animal populations (Snijders et al., 2017), and explain the structure and composition of ecological communities (Altermatt and Holyoak, 2012; Carraro et al., 2020). Rapid habitat changes are also a major threat to wildlife, as they can alter the movement patterns of individuals which may have consequences for populations (Collingham and Huntley, 2000; Todd et al., 2009). As habitat changes, individual animals can experience different spatial distributions of resources and risks, which in turn can alter the patterns of both intraspecific (Banks et al., 2007) and interspecific (Farine et al., 2015; Meise et al., 2019) interactions among individuals, as well as other processes such as dispersal patterns and gene flow (Wey et al., 2015). In population ecology, for example, changes in habitat physical configurations can reduce rates of movements among neighbouring subpopulations, which potentially reduces gene flow at the scale of meta-populations (Keller and Largiader, 2003) and impacts the persistence of populations (Frankham, 2005). Likewise, changes in habitat physical configurations could alter the transmission of information within social networks (Franz and Nunn, 2009; Whitehead and Lusseau, 2012; Barkoczi and Galesic, 2016) and other complex behavioural traits to specific social groups (Ohtsuki et al., 2007; Nowak et al., 2010; Stilwell et al., 2020). Furthermore, altered habitat physical configurations imply potential changes to the transmission dynamics of pathogens across populations (Green et al., 2006; Riley, 2007; Keeling et al., 2010; Silk et al., 2019). Our simulations show that, as animal social networks, animal habitat network structures play an important role in shaping pathogen transmission, thus highlight a fundamental link between the physical habitat environments and emergent biological processes, such as the evolutionary dynamics of cooperation (Stilwell et al., 2020) and animal culture (Somveille et al., 2018; Gruber et al., 2019).

Understanding how habitat physical configuration interacts with behavioural and/or demographic dynamics is crucial to assess how vulnerable wild animal populations – and the ecological communities that they are part of (Ryser et al., 2019) – are to the consequences of habitat changes. Our model provides the necessary first step to integrating animal movement at various spatial scales into existing quantitative frameworks. We have demonstrated that the local properties of connectivity and large-scale properties of the landscape can work together to shape population outcomes, such as the spread of pathogens. Such insights can help us to make better predictions or generate new hypotheses on how population or community structures and dynamics are shaped by the physical configurational features of habitats, and how populations or communities might respond to changing physical habitat environments.

## Acknowledgements

We thank L.M. Aplin, K.B. Beck, H.B. Brandl, J. Calatayud, M. Chimento, A.A. Maldonado-Chaparro, M. Ogino, D. Papageorgiou for their insightful comments on our manuscript, and the Max Planck Computing and Data Facility for computational support. This study was funded by the Max Planck Society, European Research Council (ERC) under the European Union’s Horizon 2020 research and innovation programme (grant agreement No. 850859), and the DFG Centre of Excellence 2117 “Centre for the Advanced Study of Collective Behaviour” under Germany’s Excellence Strategy – EXC2117 – 422037984. PH received funding from the China Scholarship Council (No.201706100183); MC was funded by a CAPES-Brazil postdoctoral fellowship (88881.170254/2018-01). We have no conflicting interest with this paper, and all authors give final approval for the publication of this paper.

## Data and code availability

Empirical data used is available from Friesen et al. (2019). The code for implementing the model is available in the R package *Animal Habitat Network* on the CRAN. The code for the simulations is available at https://github.com/ecopeng/Simulation_Code_AHN.

## Authors’ contributions

DRF and PH designed the model. PH conducted the analyses and drafted the manuscript. All authors contributed to writing and editing.

## References

Allen, B., Lippner, G., Chen, Y. T., Fotouhi, B., Momeni, N., Yau, S. T., and Nowak, M. A. (2017). Evolutionary dynamics on any population structure. Nature, 544:227–230.

Altermatt, F. and Holyoak, M. (2012). Spatial clustering of habitat structure effects patterns of community composition and diversity. Ecology, 93:1125–33.

Alther, R. and Altermatt, F. (2018). Fluvial network topology shapes communities of native and non-native amphipods. Ecosphere, 9:e02102.

Altizer, S., Nunn, C. L., Thrall, P. H., Gittleman, J. L., Antonovics, J., Cunningham, A. A., Dobson, A. P., Ezenwa, V., Jones, K. E., Pedersen, A. B., Poss, M., and Pulliam, J. R. C. (2003). Social organization and parasite risk in mammals: Integrating theory and empirical studies. Annual Review of Ecology Evolution and Systematics, 34:517–547.

Aplin, L. M., Farine, D. R., Morand-Ferron, J., and Sheldon, B. C. (2012). Social networks predict patch discovery in a wild population of songbirds. Proceedings of the Royal Society B: Biological Sciences, 279:4199–4205.

Aplin, L. M., Farine, D. R., Morand-Ferron, J., Cockburn, A., Thornton, A., and Sheldon, B. C. (2015). Experimentally induced innovations lead to persistent culture via conformity in wild birds. Nature, 518:538–541.

Armansin, N. C., Stow, A. J., Cantor, M., Leu, S. T., Klarevas-Irby, J. A., Chariton, A. A., and Farine, D. R. (2019). Social barriers in ecological landscapes: the social resistance hypothesis. Trends in Ecology and Evolution, doi: 10.1016/j.tree.2019.10.001.

Baguette, M., Blanchet, S., Legrand, D., Stevens, V. M., and Turlure, C. (2013). Individual dispersal, landscape connectivity and ecological networks. Biological Reviews, 88:31026.

Baguette, M. and Van Dyck, H. (2007). Landscape connectivity and animal behavior: functional grain as a key determinant for dispersal. Landscape Ecology, 22:1117–1129.

Banks, S. C., Piggott, M. P., Stow, A. J., and Taylor, A. C. (2007). Sex and sociality in a disconnected world: a review of the impacts of habitat fragmentation on animal social interactions. Canadian Journal of Zoology, 85:1065–1079.

Bain, G. C., Hall, M. L., and Mulder, R. A. (2014). Territory configuration moderates the frequency of extra-group mating in superb fairy-wrens. Molecular Ecology, 23:5619–5627.

Barabási, A. L., Ravasz, E., and Vicsek, T. (2001). Deterministic scale-free networks. Physica A: Statistical Mechanics and its Applications, 299:559–564.

Baranyi, G., Saura, S., Podani, J., and Jordan, F. (2011). Contribution of habitat patches to network connectivity: redundancy and uniqueness of topological indices. Ecological Indicators, 11:1301–1310.

Barkoczi, D. and Galesic, M. (2016). Social learning strategies modify the effect of network structure on group performance. Nature Communications, 7:1–8.

Barthélemy, M. (2011). Spatial networks. Physics Reports, 499:1–101.

Banks, S. C., Lindenmayer, D. B., McBurney, L., Blair, D., Knight, E. J., and Blyton, M. D. (2011). Kin selection in den sharing develops under limited availability of tree hollows for a forest marsupial. Proceedings of the Royal Society B: Biological Sciences, 278:2768–2776.

Betts, M. G., Hadley, A. S., Rodenhouse, N., and Nocera, J. J. (2008). Social information trumps vegetation structure in breeding-site selection by a migrant songbird. Proceedings of the Royal Society B: Biological Sciences, 275:2257–2263.

Bloomfield, L. S., McIntosh, T. L., and Lambin, E. F. (2020). Habitat fragmentation, livelihood behaviors, and contact between people and nonhuman primates in Africa. Landscape Ecology, 35:985–1000.

Bodin, O. and Norberg, J. (2007). A network approach for analyzing spatially structured populations in fragmented landscape. Landscape Ecology, 22:31–44.

Calizza, E., Costantini, M. L., Careddu, G., and Rossi, L. (2017). Effect of habitat degradation on competition, carrying capacity, and species assemblage stability. Ecology and Evolution, 7:5784–5796.

Cantor, M. and Farine, D. (2018). Simple foraging rules in competitive environments can generate socially structured populations. Ecology and Evolution, 8:4978–4991.

Cantor, M., Maldonado-Chaparro, A., Beck, K., Carter, G., He, P., Hillemann, F., Klarevas-Irby, J., Lang, S., Ogino, M., Papageorgiou, D., Prox, L., and Farine, D. (2019). Animal social networks: revealing the causes and implications of social structure in ecology and evolution. EcoEvoRxiv, doi: 10.32942/osf.io/m62gb.

Carraro, L., Bertuzzo, E., Fronhofer, E., Furrer, R., Gounand, I., Rinaldo, A., and Altermatt, F. (2020). Generation and application of river network analogues for use in ecology and evolution. Ecology and Evolution, doi: 10.1002/ece3.6479.

Chubaty, A., Galpern, P., and Doctolero, S. (2020). The r toolbox grainscape for modelling and visualizing landscape connectivity using spatially explicit networks. Methods in Ecology and Evolution, 11:591–595.

Collingham, Y. C. and Huntley, B. (2000). Impacts of habitat fragmentation and patch size upon migration rates. Ecological Applications, 10:131–144.

Couzin, I. D., Krause, J., Franks, N. R., and Levin, S. A. (2005). Effective leadership and decision making in animal groups on the move. Nature, 433:513–516.

Csardi, G. and Nepusz, T. (2006). The igraph software package for complex network research. InterJournal Complex Systems, 1695:1–9.

Dall, J. and Christensen, M. (2002). Random geometric graphs. Physical Review E, 66:016121.

Doherty, T. S., Fist, C. N., and Driscoll, D. A. (2019). Animal movement varies with resource availability, landscape configuration and body size: a conceptual model and empirical example. Landscape Ecology, 34:603–614.

Emlen, S. T., and Oring, L. W. (1977). Ecology, sexual selection, and the evolution of mating systems. Science, 197:215–223.

Erdős, P. and Rényi, A. (1960). On the evolution of random graphs. Bulletin of the International Statistical Institute, 38:343–347.

Fahrig, L. (2007). Non-optimal animal movement in human-altered landscapes. Functional Ecology, 21:1003–1015.

Fahrig, L. and Merriam, G. (1985). Habitat patch connectivity and population survival. Ecology, 66:1762–1768.

Fall, A., Fortin, M. J., Manseau, M., and O’Brien, D. (2007). Spatial graphs: principles and applications for habitat connectivity. Ecosystems, 10:448–461.

Farine, D. R. (2020). Structural trade-offs can predict rewiring in shrinking social networks. Journal of Animal Ecology, in press, doi: 10.1111/1365-2656.13140.

Farine, D. R., Aplin, L. M., Sheldon, B. C., and Hoppitt, W. (2015). Interspecific social networks promote information transmission in wild songbirds. Proceedings of the Royal Society B-Biological Sciences, 282:20142804.

Farine, D. R. and Sheldon, B. C. (2019). Stable multi-level social structure is maintained by habitat geometry in a wild bird population. bioRxiv, doi: 10.1101/085944.

Fischer, J., and Lindenmayer, D. B. (2007). Landscape modification and habitat fragmentation: a synthesis. Global Ecology and Biogeography, 16:265–280.

Frankham, R. (2005). Genetics and extinction. Biological Conservation, 126:131–140.

Franz, M. and Nunn, C. L. (2009). Network-based diffusion analysis: a new method for detecting social learning. Proceedings of the Royal Society B-Biological Sciences, 276:1829–1836.

Friesen, S. K., Martone, R., Rubidge, E., Baggio, J. A., and Ban, N. C. (2019). An approach to incorporating inferred connectivity of adult movement into marine protected area design with limited data. Ecological Applications, 29:e01890.

Galpern, P., Manseau, M., and Fall, A. (2011). Patch-based graphs of landscape connectivity: a guide to construction, analysis and application for conservation. Biological Conservation, 144:44–55.

Goodwin, B. and Fahrig, L. (2002). How does landscape structure influence landscape connectivity? Oikos, 99:552–570.

Gosling, L. M. (1991). The alternative mating strategies of male topi, *Damaliscus lunatus*. Applied Animal Behaviour Science, 29:107–119.

Granovetter, M. S. (1973). The strength of weak ties. American Journal of Sociology, 78:1360–1380.

Green, D. M., Kiss, I. Z., and Kao, R. R. (2006). Modelling the initial spread of foot-and-mouth disease through animal movements. Proceedings of the Royal Society B-Biological Sciences, 273:2729–2735.

Green, S. J., Boruff, B. J., and Grueter, C. C. (2020). From ridge tops to ravines: landscape drivers of chimpanzee ranging patterns. Animal Behaviour, 163:51–60.

Gruber, T., Luncz, L., Mörchen, J., Schuppli, C., Kendal, R., and Hockings, K. (2019). Cultural change in animals: a flexible behavioural adaptation to human disturbance. Palgrave Communications, 5:1–9.

Haddad, N. M., Brudvig, L. A., Clobert, J., Davies, K. F., Gonzalez, A., Holt, R. D.,… and Cook, W. M. (2015). Habitat fragmentation and its lasting impact on Earth’s ecosystems. Science Advances, 1:e1500052.

He, P. and Farine, D. (2019). Animalhabitatnetwork: networks characterising the physical configurations of animal habitats. R package, version 0.1.0 (https://CRAN.R-project.org/package=AnimalHabitatNetwork).

He, P., Maldonado-Chaparro, A. A., and Farine, D. R. (2019). The role of habitat configuration in shaping social structure: a gap in studies of animal social complexity. Behavioral Ecology and Sociobiology, 73:9.

Henriques-Silva, R., Lindo, Z., and Peres-Neto, P. (2013). A community of metacommunities: exploring patterns in species distributions across large geographical areas. Ecology, 94:627–639.

Ilany, A. and Akçay, E. (2016). Social inheritance can explain the structure of animal social networks. Nature Communications, 7:1–10.

Jacobson, B., Grant, J. W., and Pere-Neto, P. R. (2015). The interaction between the spatial distribution of resource patches and population density: consequences for intraspecific growth and morphology. Journal of Animal Ecology, 84:934–942.

Jacoby, D. M. P. and Freeman, R. (2016). Emerging network-based tools in movement ecology. Trends in Ecology and Evolution, 31:301–314.

Jordano, P. (2016). Chasing ecological interactions. PLoS Biology, 14:e1002559.

Kappeler, P. M. (2017). Sex roles and adult sex ratios: insights from mammalian biology and consequences for primate behaviour. Philosophical Transactions of the Royal Society B: Biological Sciences, 372:20160321.

Keeling, M. (2005). The implications of network structure for epidemic dynamics. Theoretical Population Biology, 67:1–8.

Keeling, M. J. (1999). The effects of local spatial structure on epidemiological invasions. Proceedings of the Royal Society of London. Series B: Biological Sciences, 266:859–867.

Keeling, M. J., and Eames, K. T. (2005). Networks and epidemic models. Journal of the Royal Society Interface, 2:295–307.

Keeling, M. J., Danon, L., Vernon, M. C., and House, T. A. (2010). Individual identity and movement networks for disease metapopulations. Proceedings of the National Academy of Sciences, 107:8866–8870.

Keller, I. and Largiader, C. R. (2003). Recent habitat fragmentation caused by major roads leads to reduction of gene flow and loss of genetic variability in ground beetles. Proceedings of the Royal Society of London. Series B: Biological Sciences, 270:417–423.

Kermack, W. and McKendrick, A. (1927). A contribution to the mathematical theory of epidemics. Proceedings of the Royal Society of London. Series A, Containing Papers of a Mathematical and Physical Character, 115:700–721.

Kokko, H. and Sutherland, W. J. (2001). Ecological traps in changing environments: ecological and evolutionary consequences of a behaviourally mediated Allee effect. Evolutionary Ecology Research, 3:603–610.

Kovalenko, K. E., Thomaz, S. M., and Warfe, D. M. (2012). Habitat complexity: approaches and future directions. Hydrobiologia, 685:1–17.

Laiolo, P. and Tella, J. L. (2005). Habitat fragmentation affects culture transmission: patterns of song matching in dupont’s lark. Journal of Applied Ecology, 42:1183–1193.

Laiolo, P. and Tella, J. L. (2006). Landscape bioacoustics allow detection of the effects of habitat patchiness on population structure. Ecology, 87:1203–1214.

Leu, S. T., Farine, D. R., Wey, T. W., Sih, A., and Bull, C. M. (2016). Environment modulates population social structure: experimental evidence from replicated social networks of wild lizards. Animal Behaviour, 111:23–31.

Loehle, C. (1995). Social barriers to pathogen transmission in wild animal populations. Ecology, 76:326–335.

Lookingbill, T. R., Gardner, R. H., Ferrari, J. R., and Keller, C. E. (2010). Combining a dispersal model with network theory to assess habitat connectivity. Ecological Applications, 20:427–41.

Lovett, G., Turner, M., Jones, C., and Weathers, K. (2007). Ecosystem function in heterogeneous landscapes. Springer, New York, NY.

Marini, L., Bartomeus, I., Rader, R., and Lami, F. (2019). Species-habitat networks: a tool to improve landscape management for conservation. Journal of Applied Ecology, 56:923–928.

McDiarmid, C., Steger, A., and Welsh, D. (2005). Random planar graphs. Journal of Combinatorial Theory Series B, 93:187–205.

McRae, B., Dickson, B., Keitt, T., and Shah, V. (2008). Using circuit theory to model connectivity in ecology, evolution, and conservation. Ecology, 89:2712–2724.

Meise, K., Franks, D. W., and Bro-Jorgensen, J. (2019). Using social network analysis of mixed-species groups in african savannah herbivores to assess how community structure responds to environmental change. Philosophical Transactions of the Royal Society B-Biological Sciences, 374:20190009.

Minor, E. S. and Urban, D. L. (2008). A graph-theory framework for evaluating landscape connectivity and conservation planning. Conservation Biology, 22:297–307.

Montiglio, P. O., McGlothlin, J. W., and Farine, D. R. (2018). Social structure modulates the evolutionary consequences of social plasticity: A social network perspective on interacting phenotypes. Ecology and Evolution, 8:1451–1464.

Mori, K., and Saito, Y. (2005). Variation in social behavior within a spider mite genus, *Stigmaeopsis* (Acari: Tetranychidae). Behavioral Ecology, 16:232–238.

Mourier, J., Vercelloni, J., and Planes, S. (2012). Evidence of social communities in a spatially structured network of a free-ranging shark species. Animal Behaviour, 83:389–401.

Naka, L. N., and Brumfield, R. T. (2018). The dual role of Amazonian rivers in the generation and maintenance of avian diversity. Science Advances, 4:eaar8575.

Nandini, S., Keerthipriya, P., and Vidya, T. N. C. (2017). Seasonal variation in female Asian elephant social structure in Nagarahole-Bandipur, southern India. Animal Behaviour, 134:135–145.

Nathan, R., Getz, W., Revilla, E., Holyoak, M., Kadmon, R., Saltz, D., and Smouse, P. (2008). A movement ecology paradigm for unifying organismal movement research. Proceedings of the National Academy of Sciences, 105:19052–19059.

Nowak, M. A., Tarnita, C. E., and Antal, T. (2010). Evolutionary dynamics in structured populations. Philosophical Transactions of the Royal Society B-Biological Sciences, 365:19–30.

Ohtsuki, H., Pacheco, J. M., and Nowak, M. A. (2007). Evolutionary graph theory: breaking the symmetry between interaction and replacement. Journal of Theoretical Biology, 246:681–694.

Pasquaretta, C., Dubois, T., Gómez-Moracho, T., Perilhon, V., Le Loc’h, G., Heeb, P., and Lihoreau, M. (2020). Analysis of temporal patterns in animal movement networks. Methods in Ecology and Evolution, in press, doi: 10.1111/2041-210X.13364.

Penrose, M. (2003). Random geometric graphs. Oxford Studies in Probability (Book 5). Oxford University Press, New York.

Perna, A. and Theraulaz, G. (2017). When social behaviour is moulded in clay: on growth and form of social insect nests. Journal of Experimental Biology, 220:83–91.

Plitzko, S. J. and Drossel, B. (2015). The effect of dispersal between patches on the stability of large trophic food webs. Theoretical Ecology, 8:233–244.

Poli, C., Hightower, J., and Fletcher Jr., R. (2020). Validating network connectivity with observed movement in experimental landscapes undergoing habitat destruction. Journal of Applied Ecology, in press, doi: 10.1111/1365-2664.13624.

Prado, F., Sheih, A., West, J., and Kerr, B. (2009). Coevolutionary cycling of host sociality and pathogen virulence in contact networks. Journal of Theoretical Biology, 261:561–569.

Prehn, S. G., Laesser, B. E., Clausen, C. G., Jønck, K., Dabelsteen, T., and Brask, J. B. (2019). Seasonal variation and stability across years in a social network of wild giraffe. Animal Behaviour, 157:95–104.

R Development Core Team (2019). R: A Language and Environment for Statistical Computing. R Foundation for Statistical Computing, Vienna, Austria (http://www.R-project.org/).

Read, J. M., and Keeling, M. J. (2003). Disease evolution on networks: the role of contact structure. Proceedings of the Royal Society of London. Series B: Biological Sciences, 270:699–708.

Riley, S. (2007). Large-scale spatial-transmission models of infectious disease. Science, 316:1298–1301.

Robertson, E., Fletcher, R., Cattau, C., Udell, B., Reichert, B., Austin, J., and Valle, D. (2018). Isolating the roles of movement and reproduction on effective connectivity alters conservation priorities for an endangered bird. Proceedings of the National Academy of Sciences, 115:8591–8596.

Ryser, R., Haussler, J., Stark, M., Brose, U., Rall, B. C., and Guill, C. (2019). The biggest losers: habitat isolation deconstructs complex food webs from top to bottom. Proceedings of the Royal Society B-Biological Sciences, 286:20191177.

Sah, P., Mann, J., and Bansal, S. (2018). Disease implications of animal social network structure: a synthesis across social systems. Journal of Animal Ecology, 87:546–558.

Silk, M. J., Hodgson, D. J., Rozins, C., Croft, D. P., Delahay, R. J., Boots, M., and McDonald, R. A. (2019). Integrating social behaviour, demography and disease dynamics in network models: applications to disease management in declining wildlife populations. Philosophical Transactions of the Royal Society B-Biological Sciences, 374:20180211.

Snijders, L., Blumstein, D. T., Stanley, C. R., and Franks, D. W. (2017). Animal social network theory can help wildlife conservation. Trends in Ecology and Evolution, 32:567–577.

Somveille, M., Firth, J. A., Aplin, L. M., Farine, D. R., Sheldon, B. C., and Thompson, R. N. (2018). Movement and conformity interact to establish local behavioural traditions in animal populations. PLoS Computational Biology, 14:e1006647.

Spear, S., Cushman, S., and McRae, B. (2015). Resistance surface modelling in landscape genetics. Landscape Genetics, 129–148, Wiley, London, UK.

Spiegel, O., Leu, S. T., Sih, A., and Bull, C. M. (2016). Socially interacting or indifferent neighbours? Randomization of movement paths to tease apart social preference and spatial constraints. Methods in Ecology and Evolution, 7:971–979.

Stilwell, P., O’Brien, S., Hesse, E., Lowe, C., Gardner, A., and Buckling, A. (2020). Resource heterogeneity and the evolution of public goods cooperation. Evolution Letters, 4:155–163.

Strandburg-Peshkin, A., Farine, D. R., Couzin, I. D., and Crofoot, M. C. (2015). Shared decision-making drives collective movement in wild baboons. Science, 348:1358–1361.

Strandburg-Peshkin, A., Farine, D. R., Crofoot, M. C., and Couzin, I. D. (2017). Habitat and social factors shape individual decisions and emergent group structure during baboon collective movement. Elife, 6:e19505.

Taylor, P. D., Fahrig, L., Henein, K., and Merriam, G. (1993). Connectivity is a vital element of landscape structure. Oikos, 68:571–573.

Tildesley, M. J., House, T. A., Bruhn, M. C., Curry, R. J., O’Neil, M., Allpress, J. L.,… and Keeling, M. J. (2010). Impact of spatial clustering on disease transmission and optimal control. Proceedings of the National Academy of Sciences, 107:1041–1046.

Todd, B. D., Luhring, T. M., Rothermel, B. B., and Gibbons, J. W. (2009). Effects of forest removal on amphibian migrations: implications for habitat and landscape connectivity. Journal of Applied Ecology, 46:554–561.

Tokeshi, M. and Arakaki, S. (2012). Habitat complexity in aquatic systems: fractals and beyond. Hydrobiologia, 685:27–47.

Tuomainen, U., and Candolin, U. (2011). Behavioural responses to human-induced environmental change. Biological Reviews, 86:640–657.

Urban, D. L. and Keitt, T. (2001). Landscape connectivity: a graph-theoretic perspective. Ecology, 82:1205–1218.

Urban, D. L., Minor, E. S., Treml, E. A., and Schick, R. S. (2009). Graph models of habitat mosaics. Ecology Letters, 12:260–273.

Vaes, O., Perna, A., and Detrain, C. (2020). The effect of nest topology on spatial organization and recruitment in the red ant *Myrmica rubra*. The Science of Nature, 107:23.

Van Schaik, C. P. (1989). The ecology of social relationships among female primates. Comparative Socioecology: The Behavioural Ecology of Humans and Other Mammals, 195–218, Blackwell, Oxford, UK.

Vähä, J. P., Erkinaro, J., Niemelä, E., Primmer, C. R. (2007). Life-history and habitat features influence the within-river genetic structure of Atlantic salmon. Molecular Ecology, 16:2638–2654.

Watts, D. J. and Strogatz, S. H. (1998). Collective dynamics of ‘small-world’ networks. Nature, 393:440–442.

Wey, T. W., Spiegel, O., Montiglio, P. O., and Mabry, K. E. (2015). Natal dispersal in a social landscape: Considering individual behavioral phenotypes and social environment in dispersal ecology. Current Zoology, 61:543–556.

Whitehead, H. and Kahn, B. (1992). Temporal and geographic variation in the social structure of female sperm whales. Canadian Journal of Zoology, 70:2145–2149.

Whitehead, H. and Lusseau, D. (2012). Animal social networks as substrate for cultural behavioural diversity. Journal of Theoretical Biology, 294:19–28.

White, L. A., Forester, J. D., and Craft, M. E. (2018). Disease outbreak thresholds emerge from interactions between movement behavior, landscape structure, and epidemiology. Proceedings of the National Academy of Sciences, 115:7374–7379.

Wilkinson, D. A., Marshall, J. C., French, N. P., and Hayman, D. T. (2018). Habitat fragmentation, biodiversity loss and the risk of novel infectious disease emergence. Journal of the Royal Society Interface, 15:20180403.

Wilson, D. S. (1975). A theory of group selection. Proceedings of the National Academy of Sciences, 72:143–146.

Wilson, M. C., Chen, X. Y., Corlett, R. T., Didham, R. K., Ding, P., Holt, R. D., Holyoak, M., Hu, G., Hughes, A. C., Jiang, L., Laurance, W. F., Liu, J. J., Pimm, S. L., Robinson, S. K., Russo, S. E., Si, X. F., Wilcove, D. S., Wu, J. G., and Yu, M. J. (2016). Habitat fragmentation and biodiversity conservation: key findings and future challenges. Landscape Ecology, 31:219–227.

Ziolkowska, E., Ostapowicz, K., Radeloff, V. C., and Kuemmerle, T. (2014). Effects of different matrix representations and connectivity measures on habitat network assessments. Landscape Ecology, 29:1551–1570.

