## Supplemental Figures for "The role of habitat configuration in shaping animal population processes: a framework to generate quantitative predictions"

PH: 0000-0002-7176-701X
POM: 0000-0002-1313-9410
MS: 0000-0002-6868-5080
MC: 0000-0002-0019-5106
DRF: 0000-0003-2208-7613

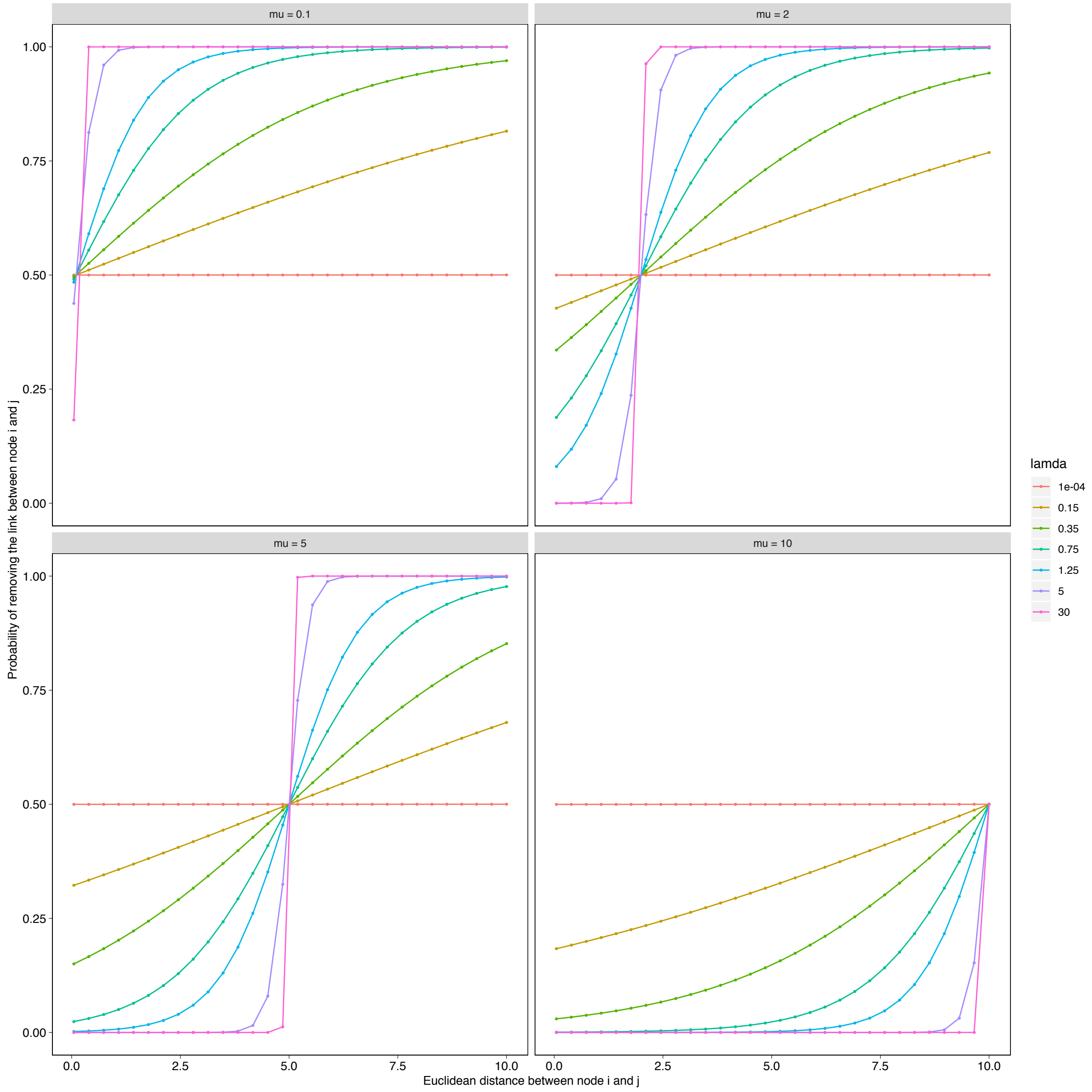


**Fig. A1** The effects of the parameter $\lambda$ (lamda) and $\mu$ (mu) on the link filtering out probability function $P(D_{ij}) = [1 + \exp(-\lambda(D_{ij} - \mu))]^{-1}$ in the AHN model. As $\lambda$ increases, the Euclidian distance between habitat components has an increasing effect on the probability of removing links from the initial complete network, while as $\mu$ increases, the probability decreases.

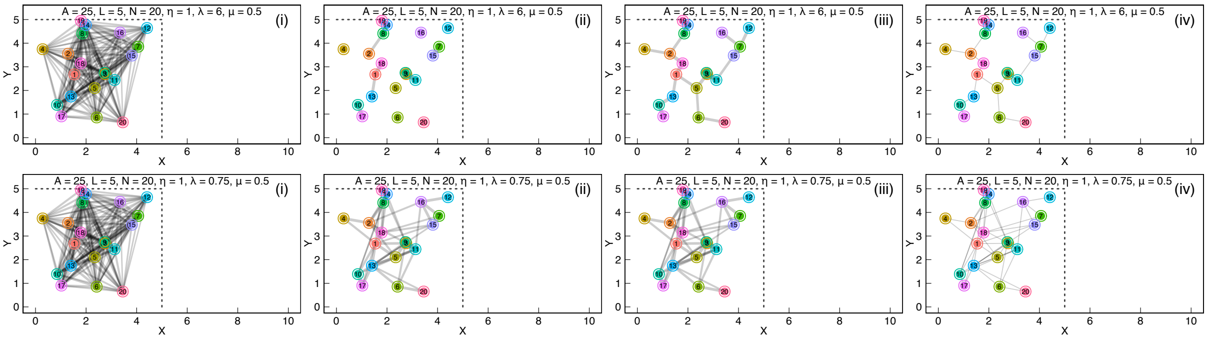


**Fig. A2** The effects of the parameter $\lambda$ in the model on the structures of the resulting animal habitat networks. As $\lambda$ decreases, the density of the habitat network tends to increase.

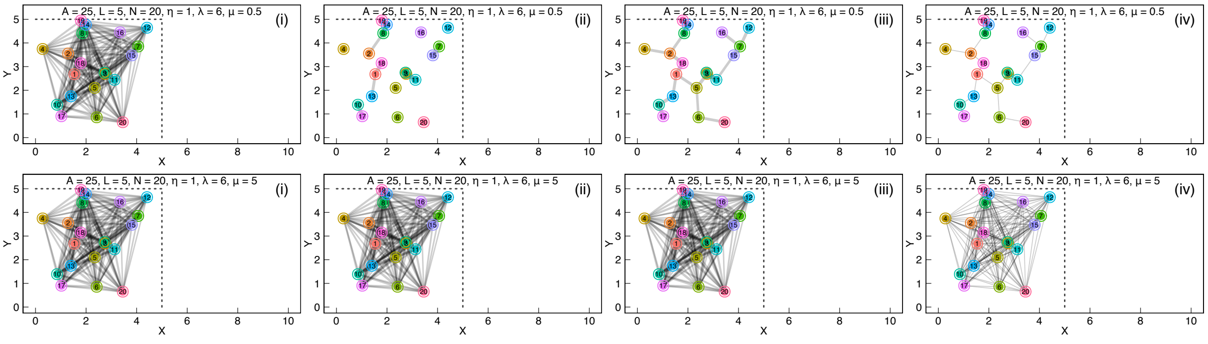


**Fig. A3** The effects of the parameter $\mu$ in the model on the structures of the resulting animal habitat networks. As $\mu$ increases, the habitat network tends to be more densely connected (less links will be removed from the initial complete network).

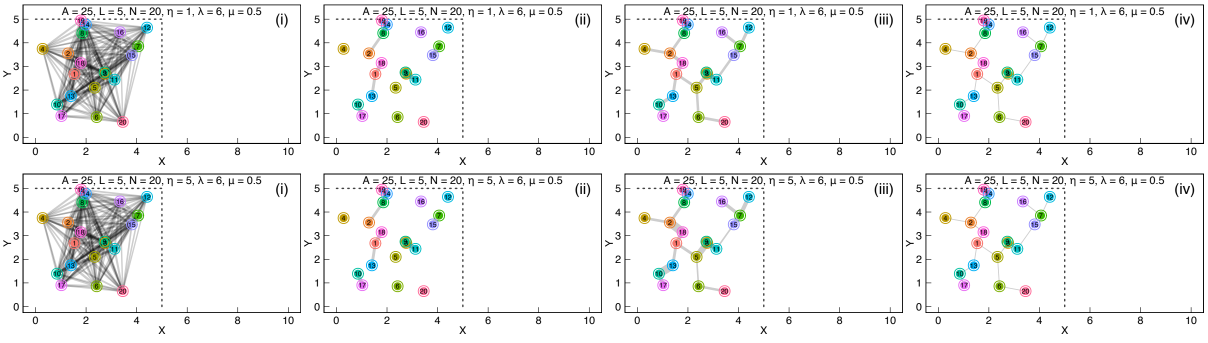


**Fig. A4** The effects of the parameter $\eta$ in the model on the structures of the resulting animal habitat networks. An increasing $\eta$ has an increasing effect on the weights of rewiring links.

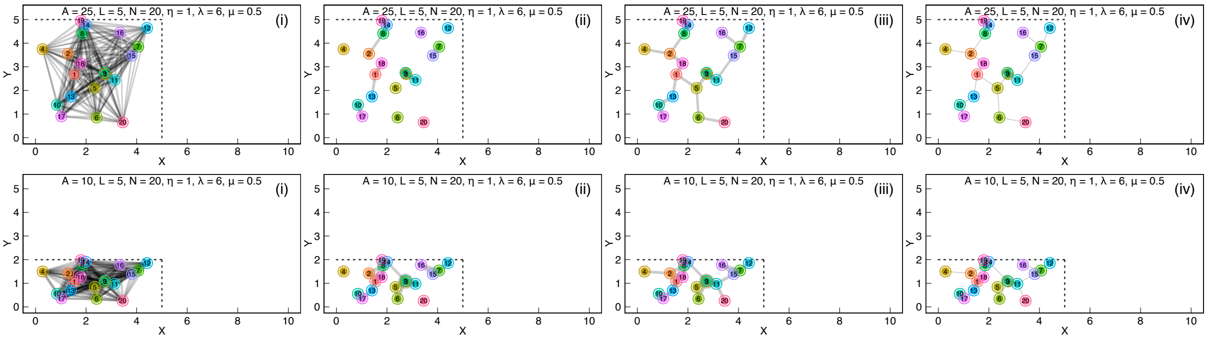


**Fig. A5** The effects of the parameter $A$ in the model on the structures of the resulting animal habitat networks. As $A$ decreases, the spatial extent at which the habitat network is defined decreases and the network tend to be more densely connected (due to a decrease in the Euclidean distances between habitat components).

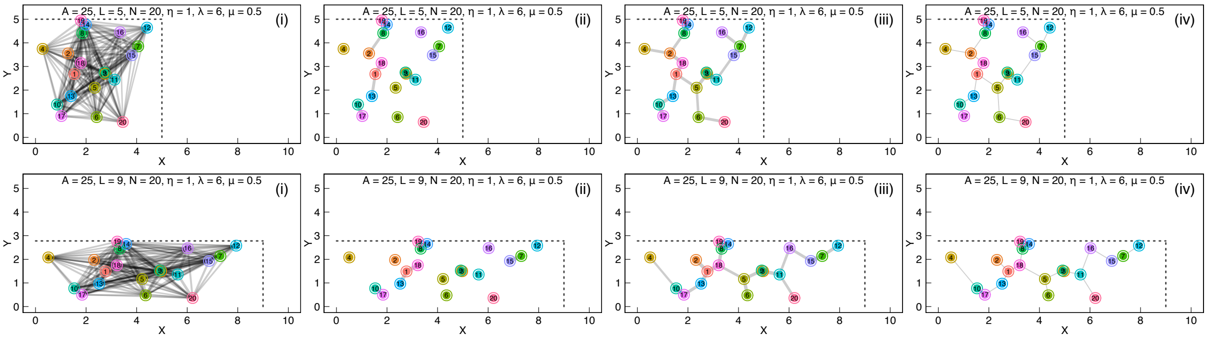


**Fig. A6** The effects of the parameter $L$ in the model on the structures of the resulting animal habitat networks. As $L$ increases, the linearity of the habitat network increases.

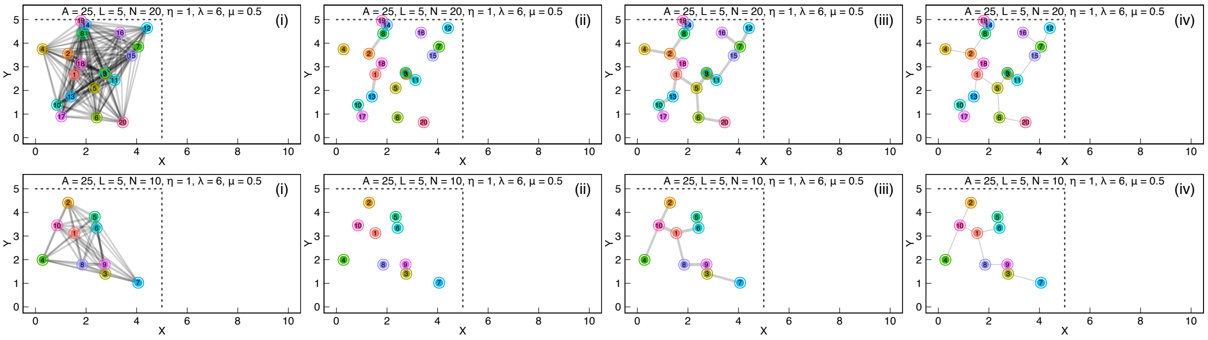


**Fig. A7** The effects of the parameter $N$ in the model on the structures of the resulting animal habitat networks. As $N$ decreases, the size and configurational complexity of the network decrease, while the spatial resolution at which the habitat network is defined increases.
